# Cancer cell-derived extracellular matrix promotes differentiation of fibroblasts into cancer-associated fibroblasts

**DOI:** 10.1101/2024.04.15.589578

**Authors:** Eyup Yondem, Devrim Pesen-Okvur

## Abstract

Breast cancer is the most common cancer and the leading cause of cancer-related mortality in women. In addition to cancer cells, the bulk of a breast tumor comprises a range of stromal cell types, including fibroblasts. Cancer-associated fibroblasts (CAF) are crucial players in the tumor microenvironment; however, the process by which fibroblasts differentiate into CAFs is not fully understood. Extracellular matrix (ECM) is known to modulate cell phenotypes. Decellularized ECM (dECM) is a useful tool for studying *in-vitro* cell-ECM interactions. Yet, whether cancer cell-derived ECM (ccECM) has a role in CAF formation is not known. Here, we optimized the culture duration (5 days) and the extraction method (freeze-thaw) for obtaining ccECM. We confirmed the presence of ccECM using coomassie blue staining and scanning electron microscopy. We showed that ccECM contained fibronectin and laminin using immunofluorescence staining. In addition, we showed that the presence of ccECM but not glass surface or TGFβ promoted the initial adhesion of fibroblasts, as expected. Finally, using quantitative immunofluorescence microscopy, we demonstrated that in contrast to fibroblasts cultured on glass surfaces in the presence and absence of TGFβ, fibroblasts cultured on ccECM showed increased expression of CAF markers vimentin (2.8 fold), FAP (3.4 fold) and PDGFR β (1.8 fold), but not FSP1/s100A4. Overall, our results indicate that ccECM promotes the differentiation of fibroblasts into CAFs.

## Introduction

The tumor microenvironment (TME) comprises both cellular and non-cellular components. It harbors cancer cells in addition to other stromal cell types, such as fibroblasts, which constitute approximately 80% of the stroma and are involved in nearly every step of tumor development in breast cancer^1^. Cancer-associated fibroblasts, also known as CAFs, are fibroblasts found in the tumor stroma that are important in cancer progression^2, 3^. The special ability of CAFs to produce highly reactive stroma facilitates extracellular matrix (ECM) turnover by increasing ECM protein synthesis and degradation by matrix-metallo-proteinases (MMPs)^4^ ^5^. Similar to cancer cells, CAF populations in tumor stroma are significantly diverse rather than homogeneous^6^. TGFβ appears as one of the strongest inducers of resident fibroblasts^7, 8^. However, it may also direct epithelial and endothelial cells to CAF differentiation over epithelial-mesenchymal transition (EMT) and endothelial-mesenchymal transition (EndMT), respectively^9–12^. It has been shown that co-culture with cancer cells can induce CAF differentiation via paracrine signaling^13, 14^. In addition, cancer cell conditioned medium (CM) could initiate fibroblast differentiation to CAFs^15^. However, it is not known whether cancer cell-derived extracellular matrix can promote differentiation of fibroblasts into CAFs.

Dynamic reciprocity defining the reciprocal connection between ECM and cells indicates that ECM can be an invaluable target for cancer research^16^. The ECM is one of the hallmarks of cancer as it is highly dysregulated and changes throughout tumor growth. These alterations encompass biochemical and biophysical changes, which in turn can impact intracellular signaling networks and change a wide range of cellular functions, such as adhesion, cell migration, invasion, proliferation, survival, epithelial-mesenchymal transition (EMT)^17–20^ .

Since ECM proteins are able to induce several phenotypes via outside-to-inside signaling, decellularized ECM (dECM) can be used as a significant tool^21^. It has been demonstrated that dECM could change cell activity through its mechanobiological ability to transmit physicochemical signals and biological functionality^22–25^. There are several studies using co-culture of cancer cells and fibroblasts or using CM from cancer cells, yet, to the best of our knowledge, the effect of cancer cell derived ECM (ccECM) on fibroblast differentiation has not been investigated.

Here, we show that decellularized ECM can be obtained from cancer cells and the resulting ccECM promotes differentiation of fibroblasts into CAFs.

## Results

### Cancer cell derived ECM (ccECM) was efficiently obtained via freeze-thaw decellularization but not chemical extraction

To reveal the role of cancer cell derived ECM (ccECM) in the differentiation of fibroblasts to CAFs, we used three different culture conditions (Fig. 1): Fibroblasts were cultured 1) on glass surfaces in standard cell culture medium, 2) on glass surfaces in standard cell culture medium supplemented with TGFβ, 3) on ccECM in standard cell culture medium.

**Figure 1.**
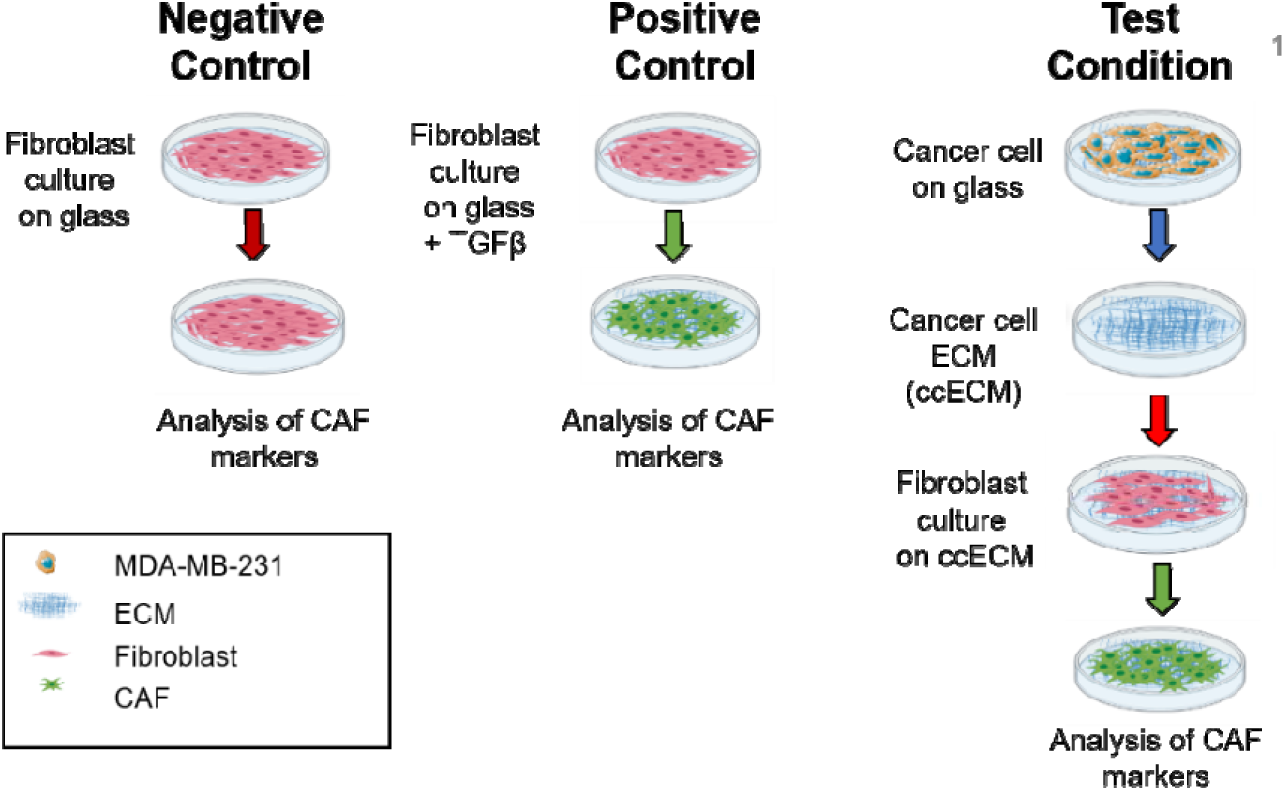
Experimental workflow. Fibroblasts were cultured on glass surface in the absence and presence of TGFβ, as negative and positive control conditions, respectively. MDA-MB-231 cells were cultured on glass surfaces, after decellurization, cancer cell ECM (ccECM) was obtained. Fibroblasts were cultured on ccECM. CAF markers were analyzed for all conditions.

To show that MDA-MB-231, a triple negative breast cancer cell line, is able to deposit ECM that can be extracted for further experiments, we compared two distinct protocols: chemical extraction and freeze-thaw decellularization on cells cultured on coverslips for 3, 5, and 7 days. Both methods resulted in uneven granular structures, more for cultures of 5 and 7 days than 3 days, as observed by light microscopy and commassie blue staining (Fig. 2A, 2B, Fig. S1). To confirm that the observed structures were indeed ECM depositions, we first used scanning electron microscopy (SEM) for cultures of 5 and 7 days (Fig. 2C). SEM images showed that surfaces prepared by freeze-thaw decellularization exhibited much more pronounced ECM-like fibrillar mesh networks than those prepared by chemical extraction on day 5. In addition, ECM-like structures were remarkably diminished on day 7, suggesting possible degradation or loss of stability over long term culture. In contrast to previously published techniques for decellularization using chemical extraction, we showed that the freeze-thaw decellularization approach was more effective for obtaining ECM produced from MDA-MB-231 cells probably because chemical extraction, which is a harsher method than freeze-thaw, has been used mostly for cells that are producing excess amount of ECM components like fibroblasts which are more active ECM producers than cancer cells which in turn tend to rather degrade ECM for invasion^26, 27^. In addition, decellularized coverslips were subjected to DNAse treatment to prevent erroneous false positive signal induction. (Fig. S2). To confirm that the observed structures were indeed ECM depositions, we finally used immunofluorescence staining for two of the most prominent ECM glycoproteins: fibronectin and laminin(Fig.2D). Fluorescence microscopy images clearly showed presence of both fibronectin and laminin where fibronectin signal was 2.59 fold higher than laminin signal. (Fig.2E, Fig.2F, n=3, ****p< 0,0001). This outcome was consistent with recent *in vivo* findings showing the ECM from TNBC contains more fibronectin than laminin.^23^

**Figure 2.**
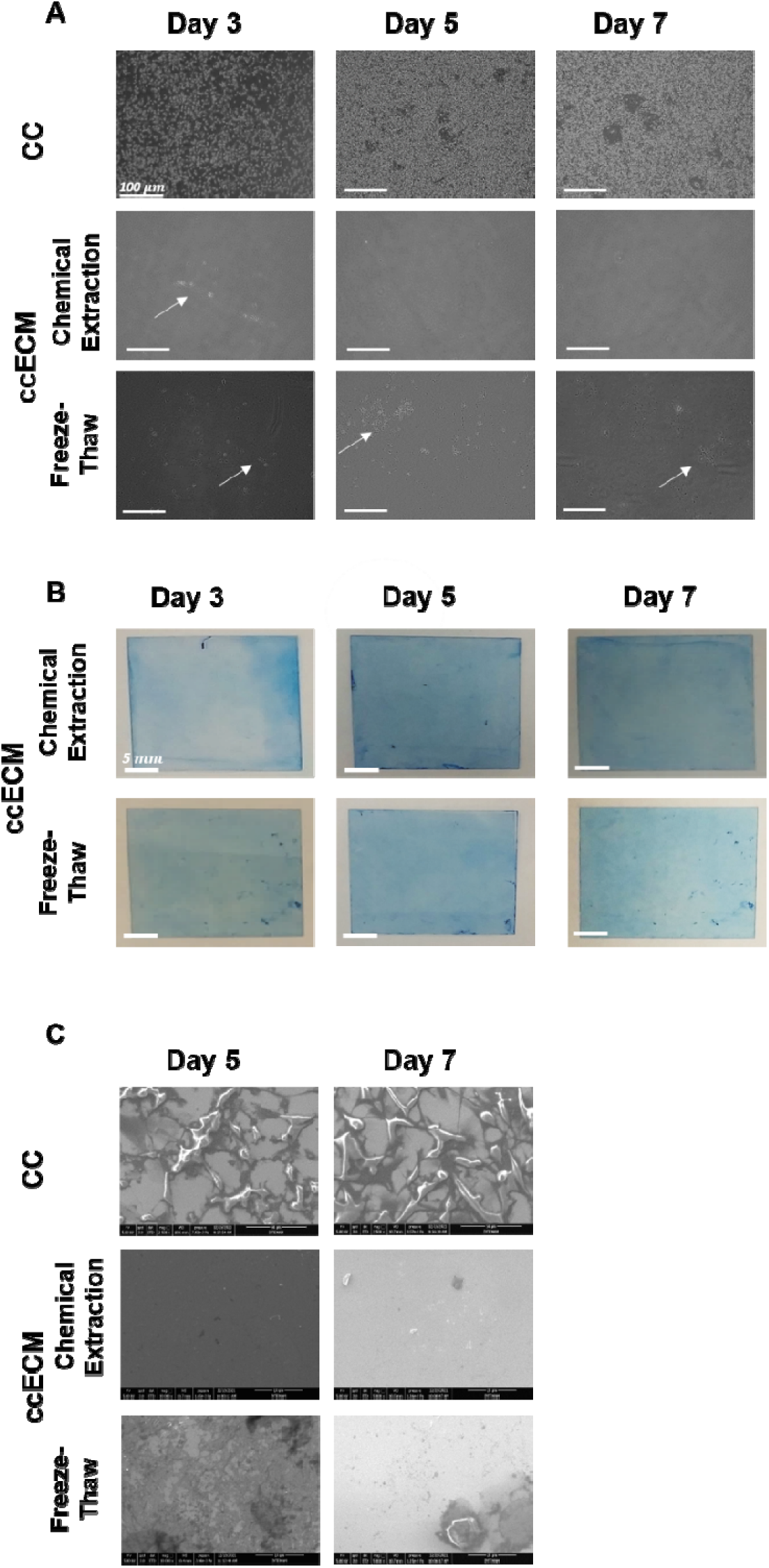
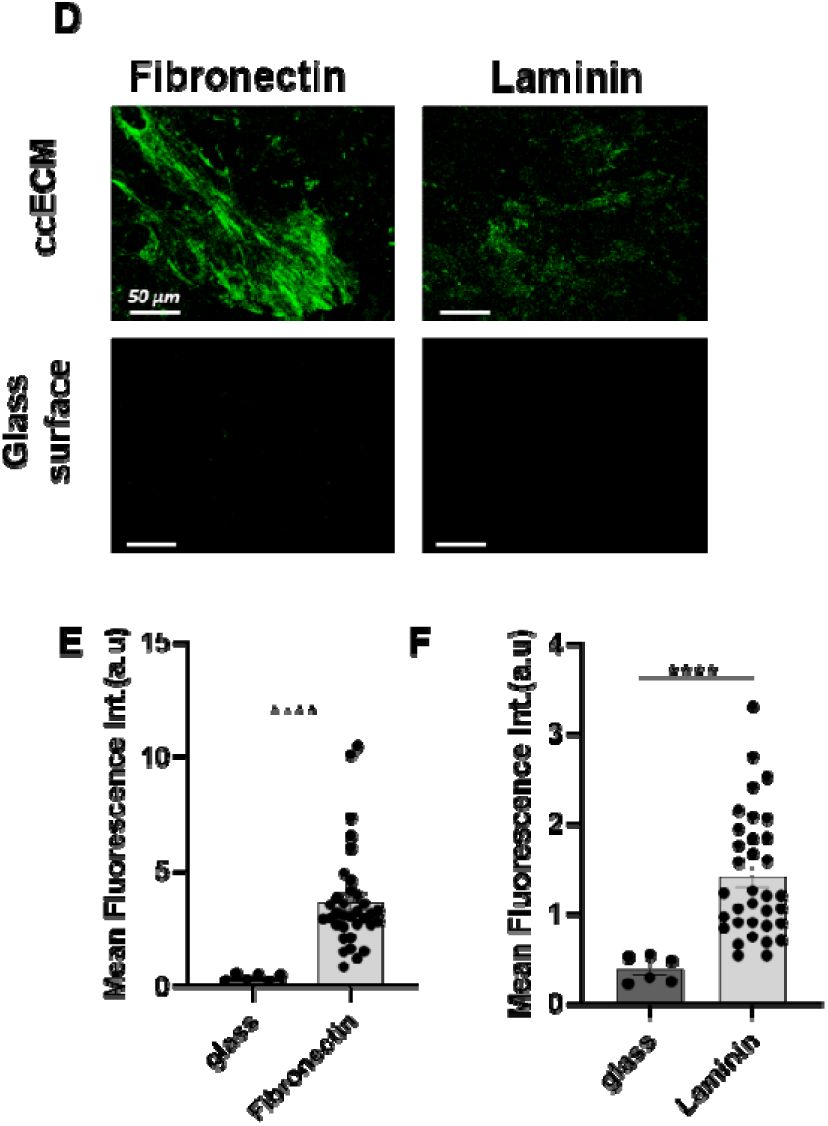
Cancer cell-derived ECM (ccECM) was efficiently obtained via freeze-thaw decellularization but not chemical extraction. A. Phase contrast images. Top panel MDA-MB-231 cells (cc) in culture, middle panel ccECM obtained using chemical extraction, bottom panel ccECM obtained using freeze-thaw. Cells were cultured for 3, 5, or 7 days. B. Coomassie Blue staining for qualitative analysis of ECM deposition top panel ccECM obtained using chemical extraction, bottom panel ccECM obtained using freeze-thaw. C. Scanning electron microscopy images. Top panel MDA-MB-231 cells (cc) in culture, middle panel ccECM obtained using chemical extraction, bottom panel ccECM obtained using freeze-thaw. D. Immunofluorescence images of fibronectin or laminin for ccECM and control glass surface. Mean fluorescence intensities were plotted for different conditions. (* p < 0.05; ** p < 0.01; *** p < 0.005; **** p < 0.0001, n=3).

### ccECM promoted initial adhesion of fibroblasts in contrast to glass surfaces in the presence and absence of TGFβ

We cultured WI-38, a lung fibroblast cell line, using the three different culture conditions as defined above. When we examined confluency of fibroblasts at the end of 48 hours by labeling the cell nuclei with DAPI, there were 2.3 and 2.1 fold more cells on the ccECM surface than the glass surface in the absence of TGFβ and the glass surface in the presence of TGFβ, respectively (Fig. 3A, n=3, ****p< 0,005). To differentiate cell adhesion from proliferation, we compared surface coverage at different time points using fibroblasts labeled with green fluorescent cell tracker (Fig.3B). At 2 hrs, fibroblasts already showed 1.3 fold more cell adhesion on the ccECM surface than the other two conditions (Fig. 3C, n=3, ***p< 0.005(TGFβ, ***p<0.005(Glass)). However, the rates of increase in confluency, determined by the slopes of the linear trend lines, were similar for all three conditions (Fig. 3D). Thus, the higher confluency in later time points was due to enhanced initial cell adhesion.

**Figure 3.**
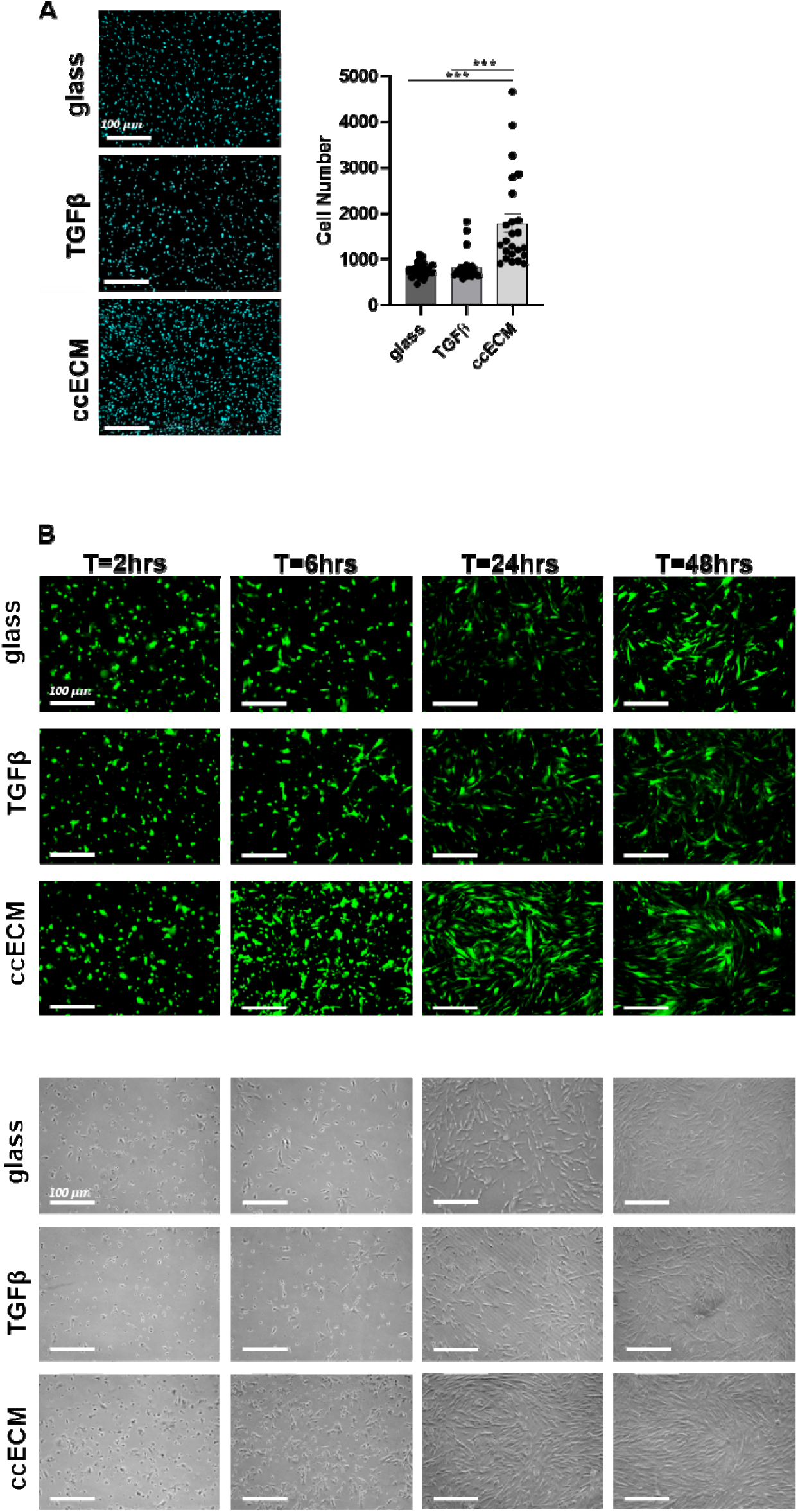
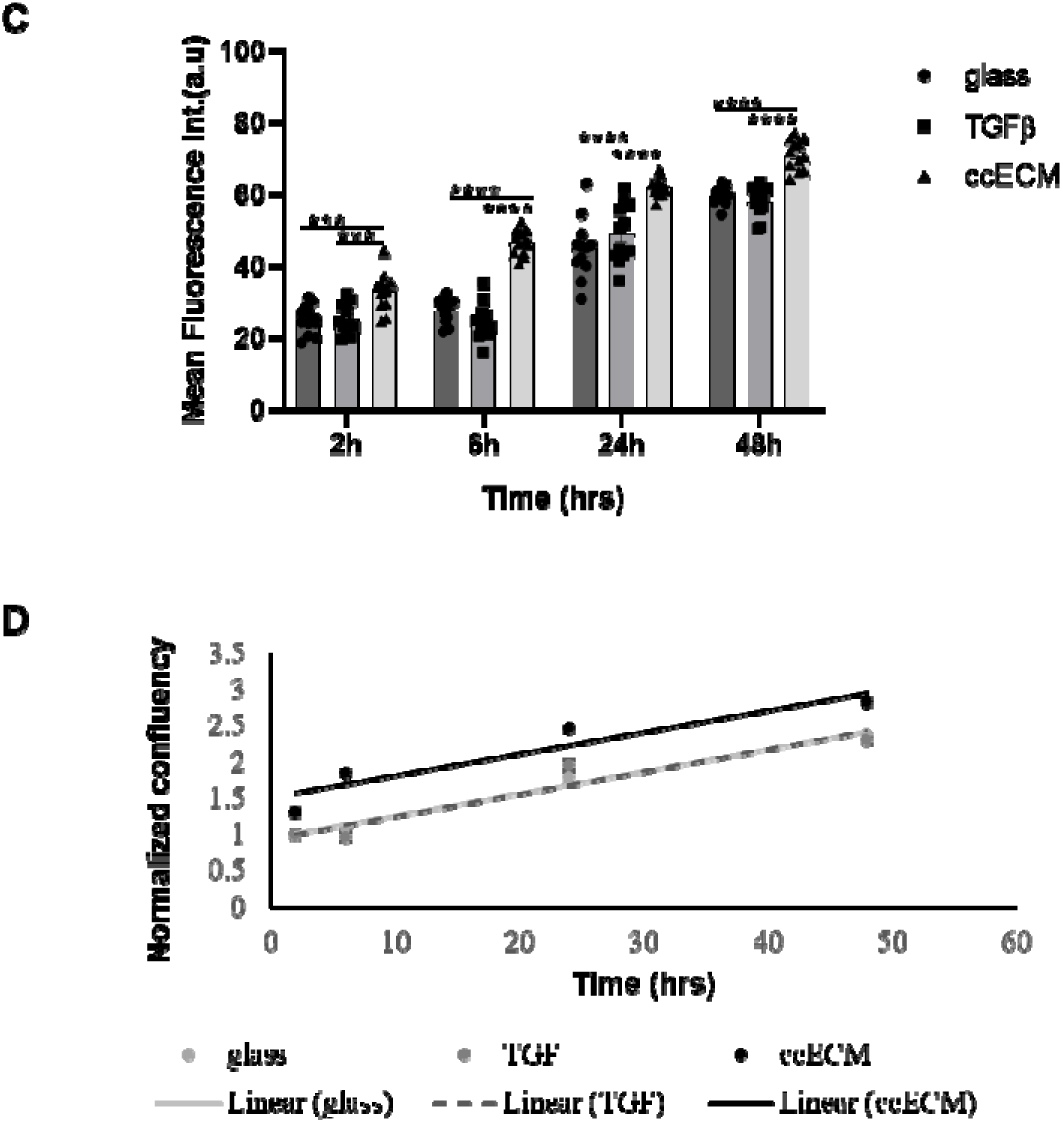
ccECM promoted initial adhesion of fibroblasts in contrast to glass surfaces in the presence and absence of TGFβ. Fluorescence images of DAPI stained cells cultured on the glass surface in the absence of TGFβ (glass), on the glass surface in the presence of TGFβ, and on the ccECM (ccECM) for 48 hrs. Mean cell numbers were plotted for different conditions (n=8). B. Fluorescence (top) and brightfield (bottom) images of Cell tracker^TM^ Green CMFDA dye-labeled fibroblasts cultured on the glass surface in the absence of TGFβ (glass), on the glass surface in the presence of TGFβ, and on the ccECM (ccECM) for 2hrs, 6hrs, 24hrs, and 48hrs. C. Mean GreenTracker^TM^ fluorescence intensity was plotted for different conditions (* p < 0.05; ** p < 0.01; *** p < 0.005; **** p < 0.0001, n=3). D. Confluency as a function of time normalized to confluency on glass surface at 2 hrs.

### ccECM enhanced expression of CAF markers as well as or more than TGFβ

We cultured WI-38 fibroblasts, using the three different culture conditions as defined above and examined the expression of CAF markers. The common cytokine TGFβ is known to stimulate differentiation of fibroblasts ^28–31^. Thus, glass surface, where we employed 10ng/ml TGFβ, acted as a positive control for fibroblast differentiation. Among the commonly used CAF markers in the literature, we tested for vimentin, FAP, PDGFRβ, and FSP-1/s100A4^32, 33^.

Vimentin is a type III intermediate filament^34^. Using immunofluorescence, we showed that vimentin expression was 2.7 fold higher on glass surfaces in the presence of TGFβ than in its absence (Fig.4A, 4B, n=3, **** p <0.0001), indicating TGFβ-induced fibroblast differentiation. In addition, vimentin expression was 2.8 fold higher on the ccECM surface than the glass surface in the absence of TGFβ and similar to the glass surface in the presence of TGFβ (Fig.4A, 4B, n=3, ****p <0.0001). Meanwhile, RT-PCR results revealed that mRNA expression levels did not show any significant differences between the three different culture conditions (Fig.4C). Vimentin expression was reported to be confluency-dependent in numerous studies such that vimentin expression rose as confluency dropped^35,36^. To clarify this point, we used different initial cell seeding densities of 5208 cells/cm^2^ (low) and 52080 cells/cm^2^ (high) (Fig.4D). Immunofluorescence results showed that regardless of confluency, vimentin expression was 2.06 (low cell density) and 2.33 (high cell density) fold higher on the ccECM surface compared to the glass surface in the absence of TGFβ (Fig.4E, * p<0.05). These results showed that ccECM can induce expression of vimentin, a CAF marker, as well as TGFβ.

**Figure 4.**
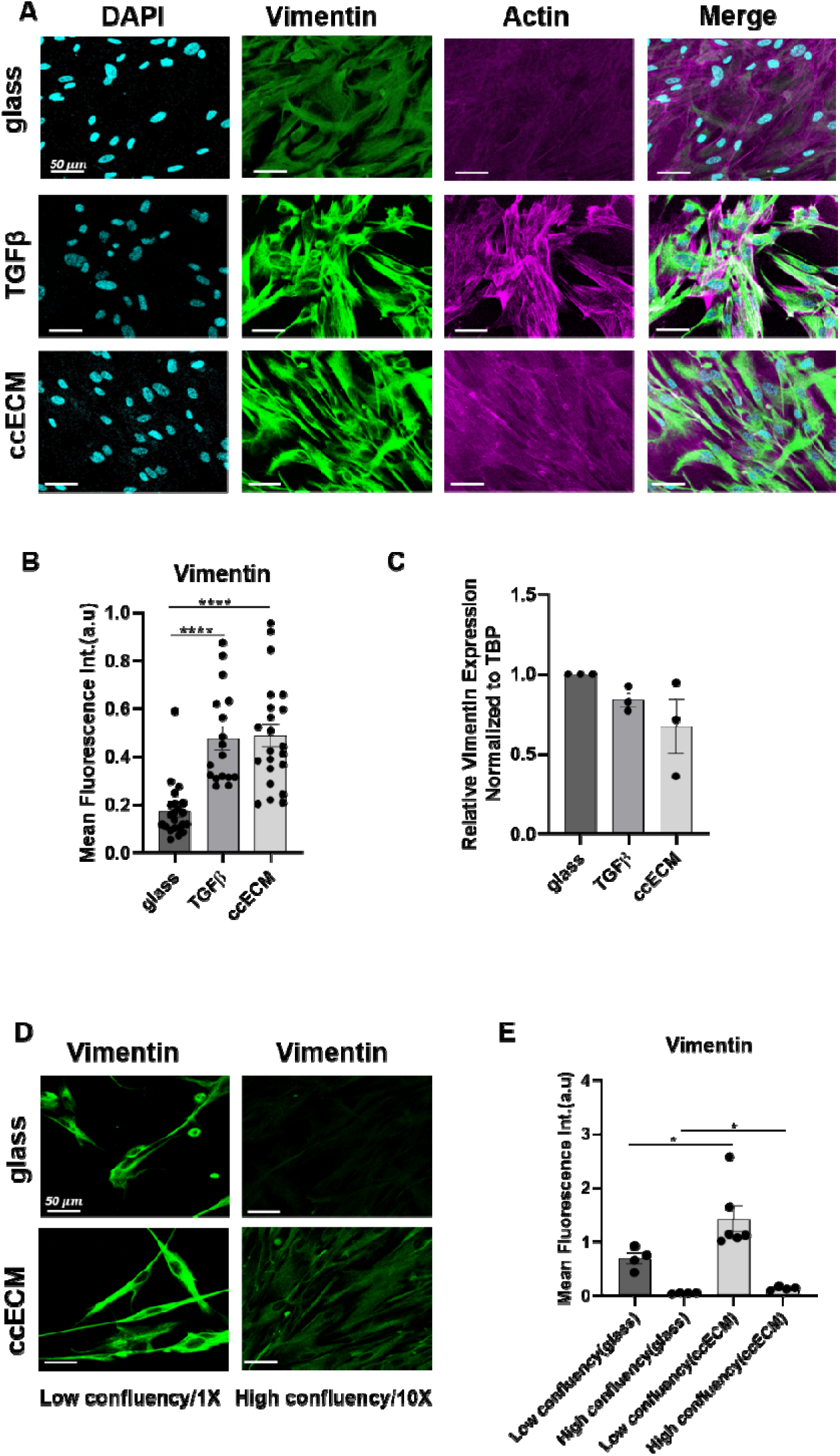
ccECM enhanced expression of CAF markers as well as or more than TGFβ: Vimentin. A. Immunofluorescence images of cells cultured on the glass surface in the absence of TGFβ (glass), on the glass surface in the presence of TGβ and on the ccECM (ccECM) for 48 hrs, stained for nucleus (DAPI) (cyan), vimentin (green), and actin (magenta). B. Quantification of vimentin intensity normalized to cell number (n=3) C. Relative mRNA expression of vimentin normalized to TBP (n = 3). D. Immunofluorescence images of low or high confluency cells cultured on the glass surface in the absence of TGFβ (glass), and on the ccECM (ccECM) for 48 hrs, stained for vimentin (green). E. Quantification of vimentin intensity normalized to cell number. Mean fluorescence intensity was plotted for different conditions (* p < 0.05; ** p < 0.01; *** p < 0.005; **** p < 0.0001).

Fibroblast Associated Protein(FAP), which is a type II serine protease present at the cell surface, is another CAF marker^37^. Using immunofluorescence, we showed that FAP expression was 3.4 and 1.9 fold higher on the ccECM surface than the glass surface in the absence of TGFβ and the glass surface in the presence of TGFβ, respectively (Fig.5A, 5B, n=2, **** p <0.0001). In addition, FAP expression was 1.7 fold higher on the glass surface in the presence of TGFβ than in the absence of TGFβ (Fig.5A, 5B, n=2, **** p <0.0001). At the RNA level, FAP expression was 2.5 and 3.7 fold higher on glass surface in the presence of TGFβ than on glass surface in the absence of TGFβ and on the ccECM surface, respectively (Fig5C, n=3, **** p< 0.00001). These results showed that ccECM can induce expression of FAP more than TGFβ at the protein level, but not the RNA level.

**Figure 5.**
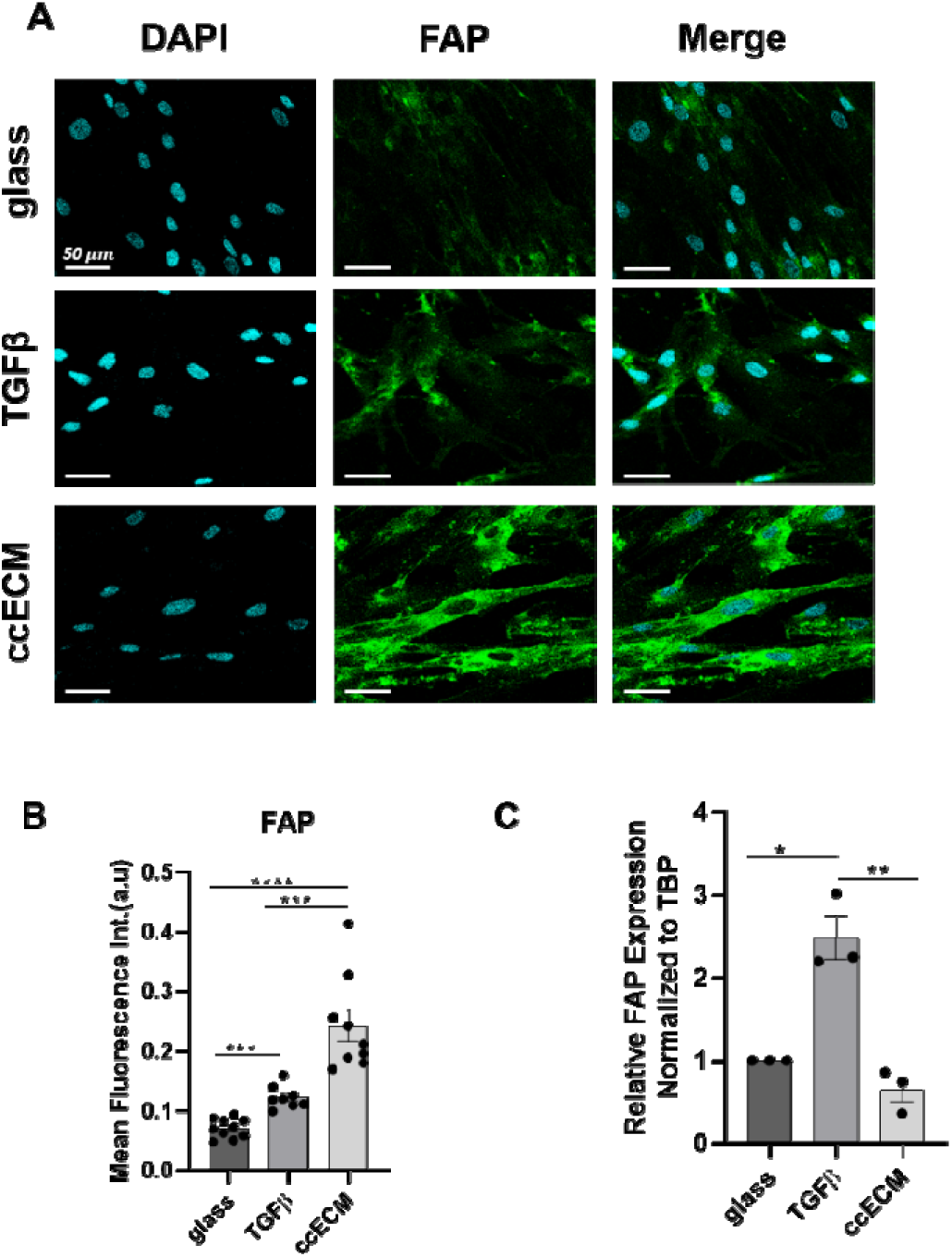
ccECM enhanced expression of CAF markers as well as or more than TGFβ: Fibroblast Associated Protein (FAP). A. Immunofluorescence images of cells cultured on the glass surface in the absence of TGFβ (glass), on the glass surface in the presence of TGFβ, and on the ccECM (ccECM) for 48 hrs, stained for nucleus (DAPI) (cyan), vimentin (green). B. Quantification of FAP intensity normalized to cell number (n=3) C. Relative mRNA expression of FAP normalized to TBP (n = 3). Mean fluorescence intensity was plotted for different conditions (* p < 0.05; ** p < 0.01; *** p < 0.005; **** p < 0.0001).

Platelet-derived growth factor receptor beta (PDGFRβ), which is a type III tyrosine-protein kinase receptor present on the cell surface^38^, is another CAF marker. Immunofluorescence imaging showed expression of PDGFRβ was 1.8 and 1.5 fold higher on the ccECM surface than the glass surface in the absence of TGFβ and the glass surface in the presence of TGFβ, respectively (Fig.6A, 6B, n=3, **** p <0.0001). In addition, PDGFRβ expression was 1.2 fold higher on the glass surface in the presence of TGFβ than in the absence of TGFβ (Fig.6A, 6B, n=3, **** p <0.0001). These results mirrored the results for FAP. Yet, at the RNA level, there were no significant differences as was the case for vimentin (Fig.6C n=3). Together, these results indicate that ccECM can induce expression of PGFRβ more than TGFβ at the protein level.

**Figure 6.**
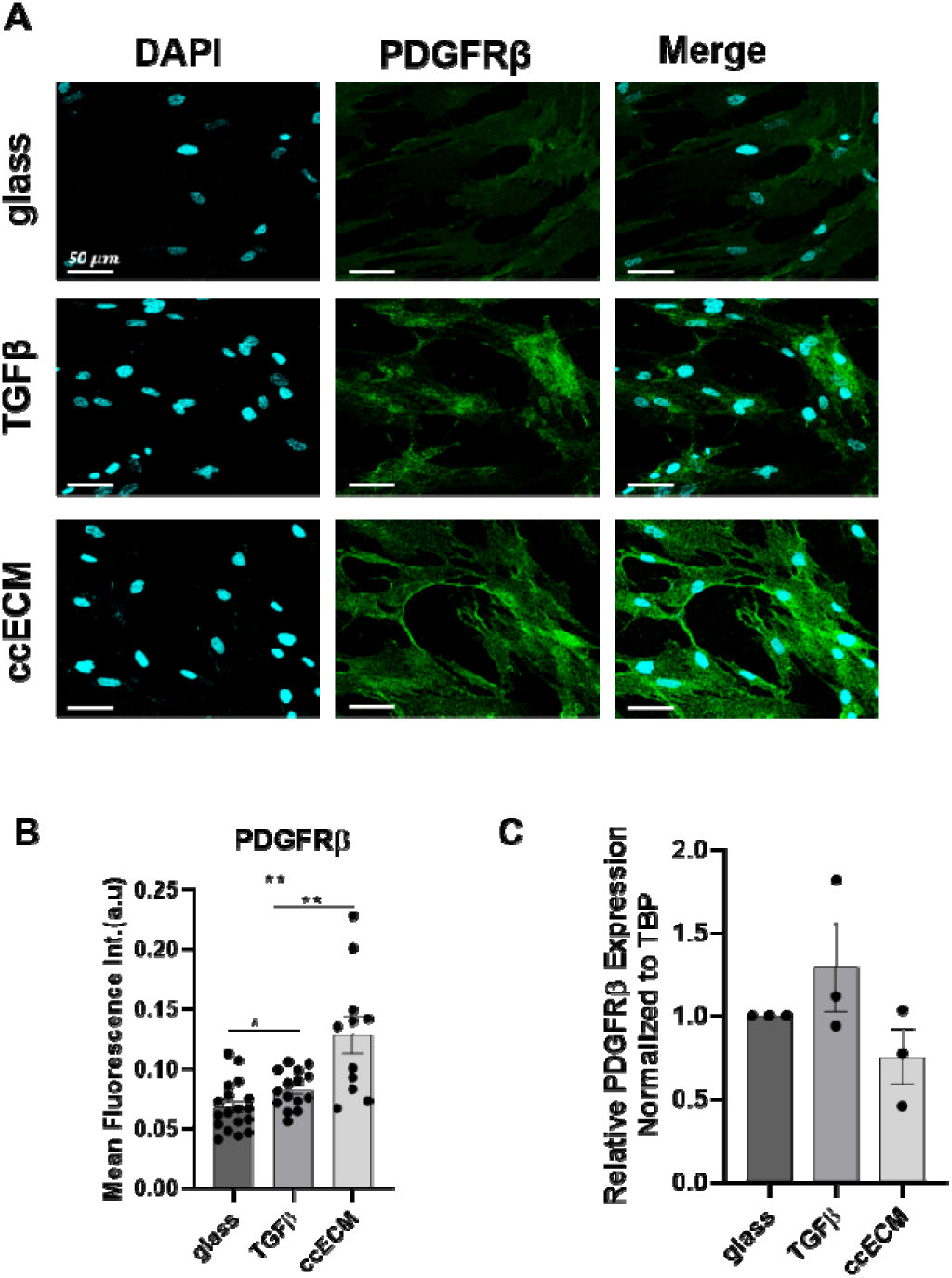
ccECM enhanced expression of CAF markers as well as or more than TGFβ: Platelet-derived growth factor receptor beta (PDGFRβ). A. Immunofluorescence images of cells cultured on the glass surface in the absence of TGFβ (glass), on the glass surface in the presence of TGFβ, and on the ccECM (ccECM) for 48 hrs, stained for nucleus (DAPI) (cyan), PDGFRβ (green). B. Quantification of PDGFRβ intensity normalized to cell number (n=3) C. Relative mRNA expression of FAP normalized to TBP (n = 3). Mean fluorescence intensity was plotted for different conditions (* p < 0.05; ** p < 0.01; *** p < 0.005; **** p < 0.0001).

Another CAF marker we investigated was FSP1/s100A4, a fibroblast-specific filament-associated protein belonging to the s100 cytoplasmic calcium-binding protein family^39^. Interestingly, at the protein level, there were no significant differences between the three different culture conditions (Fig. 7A, 7B, n=3) whereas at the RNA level, cells showed similar levels of expression on glass surface in the absence of TGFβ and on ccECM surface, which were 3.1 and 3.4 fold higher than that on glass surface in the presence of TGFβ (Fig. 7C, n=3, *** p <0.005, p<0.05).

**Figure 7.**
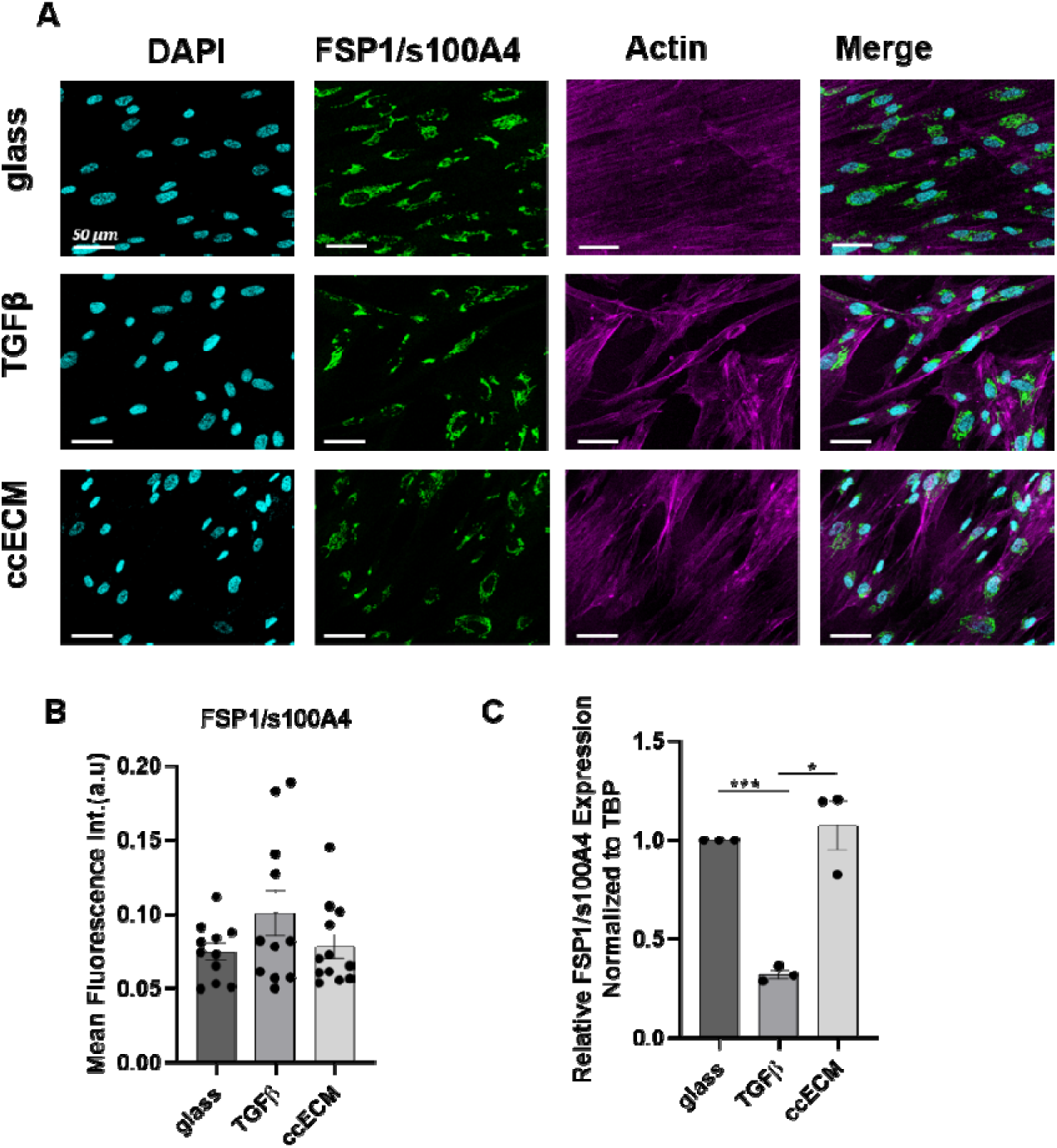
ccECM enhanced expression of CAF markers as well as or more than TGFβ: Fibroblast-specific filament-associated protein(FSP1/s100A4). A.Immunofluorescence images of cells cultured on the glass surface in the absence of TGFβ (glass), on the glass surface in the presence of TGFβ, and on the ccECM (ccECM) for 48 hrs, stained for nucleus (DAPI) (cyan), FSP1 (green), and actin(magenta). B. Quantification of FSP1 intensity normalized to cell number (n=3) C. Relative mRNA expression of FSP1 normalized to TBP (n = 3). Mean fluorescence intensity was plotted for different conditions ((* p < 0.05; ** p < 0.01; *** p < 0.005; **** p < 0.0001).

### Fibroblasts cultured on ccECM expressed lower levels of fibronectin and laminin

The results on CAF markers on the ccECM surfaces supported the presence of outside-inside signaling originating from ccECM. Thus, we tested whether another CAF-associated phenotype, modified ECM deposition, is regulated by ccECM by comparing the expression levels of the two ECM proteins, fibronectin and laminin for fibroblasts cultured in the three conditions we have used. Fibronectin expression in fibroblasts cultured on the glass surface in the presence of TGFβ was 1.5 fold higher than those on the ccECM surface (Fig. 8A, 8C n=3, **p<0.01). What is more, laminin expression was 2.3 fold higher on the glass surface in the absence of TGFβ than that on the ccECM surface and 1.9 fold higher on the glass surface in the presence of TGFβ than that on the ccECM surface (Fig. 8B, 8D, n=3, ***p< 0.005).

**Figure 8.**
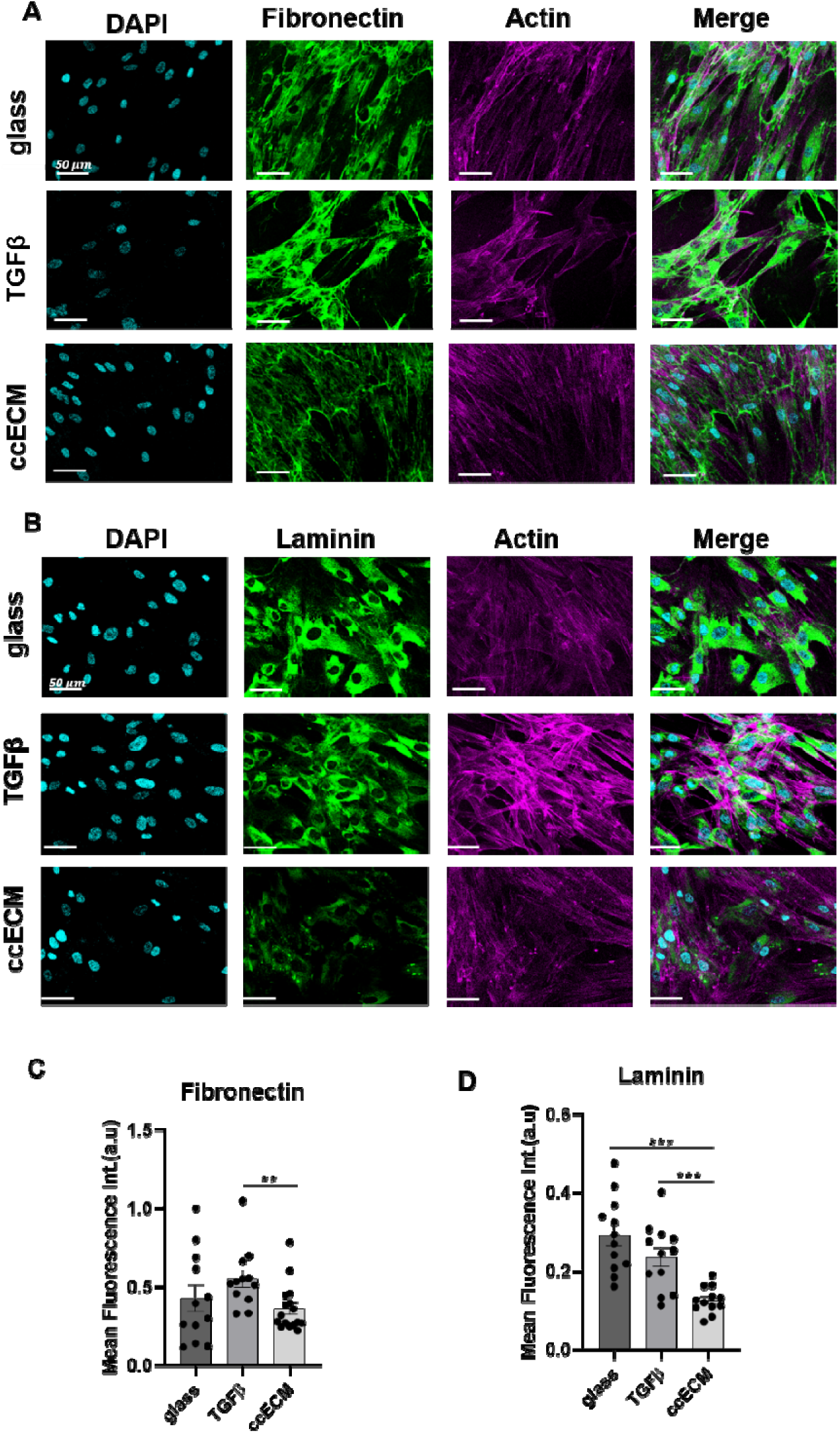
Fibroblasts cultured on ccECM expressed lower levels of fibronectin and laminin. A. Immunofluorescence images of cells cultured on the glass surface in the absence of TGFβ (glass), on the glass surface in the presence of TGFβ, and on the ccECM (ccECM) for 48 hrs, stained for nucleus (DAPI) (cyan), Fibronectin (green), and actin(magenta). B. Immunofluorescence images of cells cultured on the glass surface in the absence of TGFβ (glass), on the glass surface in the presence of TGFβ, and on the ccECM (ccECM) for 48 hrs, stained for nucleus (DAPI) (cyan), Laminin (green), and actin (magenta). C. Quantification of Fibronectin intensity normalized to cell number (n=3) D. Quantification of Laminin intensity normalized to cell number (n=3). Mean fluorescence intensity was plotted for different conditions (* p < 0.05; ** p < 0.01; *** p < 0.005; **** p < 0.0001).

## Discussion

We investigated whether decellurized ECM can be obtained from cancer cells and the resulting ccECM can promote differentiation of fibroblasts into CAFs. We showed that the freeze-thaw method was more efficient than chemical extraction for decellularization. Chemical extraction, which uses triton X-100 and ammonium hydroxide, appeared to be too strong for ECM produced by cancer cells. Verstegen et. al., previously showed that triton X-100 might decrease fibronectin and laminin on the decellularized ECM, causing the dECM structure to be disturbed^40^, consistent with our results. What is more, Hielscher et. al., noted that obtaining decellularized ECM using breast cancer cells (BCCs), notably MDA231 cells, presented difficulties. Therefore they solved this problem by coculturing MDA-MB-231 cells with fibroblasts and chemical extraction can be effective, yet, the resulting dECM was not purely from cancer cells^41^.

By analyzing expression of CAF markers at the protein and RNA levels, we examined the differentiation of the fibroblast grown on ccECM to CAFs. Vimentin, FAP, PDGFRβ, and FSP1/s100A are among the commonly used CAF markers in the literature ^42^. Notably, except for FSP1/s100A, a significant and consistent increase was observed at the protein level. Despite the fact that FSP1/s100A4 is one of the CAF markers, recent studies have shown that FSP1 expression fluctuates in the population of CAFs in diverse ways, in agreement with our results ^43–45^. By analyzing the relative mRNA expression level of CAF markers, there were two markers that showed significantly differences in fibroblasts on ccECM surface, where expression of FAP dropped and FSP1 increased compared to the glass surface in the presence of TGFβ. In addition to having growth factors, chemokines, and cytokines, dECM also contains ECM elements such as collagen, laminin, and fibronectin^46–48^. Additionally, ECM components could alter as cells develop into malignant ones, and the ECM gathers various substances in contrast to its healthy counterpart^49, 50^. As a result, fibroblasts grown on ccECM and TGF-induced fibroblasts may not necessarily express an identical set of CAF markers. Since the components in the ccECM may have an impact on how these proteins are translated, the half-life of the mRNA may differ from that of other groups, leading to enhanced protein expression but not mRNA^51^.

Despite ongoing debate regarding vimentin’s classification as a CAF marker, multiple articles demonstrated a consistent elevation in vimentin expression when fibroblasts were co-cultured with cancer cells, which coincided with the differentiation of fibroblasts into CAFs. These findings contribute to the understanding of CAF biology and emphasize the potential significance of vimentin as an important marker in the process of fibroblast differentiation into CAFs^52, 53^. Patients with vimentin-positive CAFs also had a worse prognosis and a lower overall survival rate^54^. Vimentin expression was reported to be confluency-dependent: vimentin expression increased as confluency decreased^35, 36^. Therefore, we also examined the effect of confluency. Independent of the confluency, we observed a considerable increase for vimentin.

We examined laminin and fibronectin^55^, two less obvious CAFs indicators, in addition to CAFs markers. In the context of our experiment, fibronectin on the ccECM surface remained constant as compared to the glass surface in the absence of TGFβ but not the glass surface in the presence of TGFβ. Compared to the glass surface in the presence of TGFβ, fibronectin was much lower in the fibroblast on ccECM surface. Laminin expression was also considerably reduced on the ccECM surface compared to the glass surface in the absence of TGFβ and the glass surface in the presence of TGFβ. Since ccECM already contained these ECM components in its structure, we thus reasoned that there might be a negative feedback in the case of the ccECM surface (Fig.2D).

When fibroblasts were cultured on the ccECM, they exhibited enhanced attachment and confluence to compared to the other two culture conditions (Fig. 3). Karina et. al., previously showed improved attachment for fibroblasts and CAFs on a substrate rich in fibronectin^56^. In agreement, ccECM, which was rich in fibronectin and laminin, supported initial cell adhesion.

Overall, ccECM appears to be one of the crucial mechanobiological players in fibroblast differentiation. Our results support the hypothesis that ccECM could differentiate fibroblasts into CAFs. Future research will examine the ccECM comprehensively to decipher the mechanisms underlying this transdifferentiation and whether ECM derived from other cell types can have similar effects. The innovative perspective we present is expected to contribute to a deeper understanding of cell-to-matrix communication in various health and disease contexts.

### Experimental procedures Cell culture

MDA-MB-231 (invasive triple negative) breast cancer cell line and WI-38 (lung derived) fibroblastic cell line were acquired from ATCC. MDA-MB-231 cells and WI-38 cells were maintained in DMEM (Biological Industries - BI), and MEM-ALPHA (BI) supplemented with 10% FBS(BI), 1% L-Glutamine (BI), and 1% Penicillin/Streptomycin (P/S) (BI), respectively. All cells were cultured at 37°C in a humidified incubator with 5% CO2. WI-38 cells were stained with 5μM/ml Cell Tracker^TM^, Green CMFDA Dye (Invitrogen, C2925) according to manufacturer’s protocol.

### Isolation of Cell-Derived Extracellular Matrix Chemical extraction

ccECM extraction from MDA-MB-231 cells was performed as previously described^27^. Briefly, breast cancer cells were seeded at 5×10^4^ cells/cm^2^ onto glass coverslips for 3, 5, and 7 days, and half of the culture medium was replaced every two days. The coverslips were maintained within the humidified atmosphere at 5% CO2 and 37°C. Once the culture time ended, coverslips were exposed to the extraction buffer containing 20mM NH4OH (Sigma), 0.05% Triton X-100/ 1X Phosphate Buffer Saline (PBS) until cells were completely removed from the surface of the coverslips. During detachment, coverslips were observed under light microscopy to ensure cells washed away completely.

### Freeze-Thaw

MDA-MB-231 cells were seeded onto coverslips and cultured similar to the extraction buffer process. However, once the culture time finished, the coverslips were exposed to rapid freezing at -80°C for 35 minutes, and then thawing with warm (37°C) 1X PBS wash to remove the cell debris leaving behind ccECM. If the cells did not detach at the first cycle, coverslips were processed for one more cycle of freeze-thaw. Unless otherwise indicated, coverslips were exposed to only one freeze-thaw cycle.

### Qualitative Determination of ECM

Coomassie blue staining was used to detect presence of ccECM proteins. 5×10^4^ cells/cm^2^ MDA-MB-231 cells were cultivated on coverslips for 3, 5, and 7 days. Then, coverslips were decellularized either by chemical extraction or freeze-thaw. Decellularized coverslips were shaken for up to 30 minutes while being incubated with a 0.1% Coomassie Blue solution that included 70% H_2_O, 25% methanol, and 5% acetic acid. The coverslips were then dipped once in H_2_O and air-dried.

### Attachment and Proliferation Assay

Fibroblast cells labeled with Cell Tracker^TM^, Green CMFDA Dye were cultured under three conditions: cells on ccECM-containing coverslips, ccECM-free coverslips with TGFβ treatment, and ccECM-free coverslips without TGFβ treatment. Then images were captured using fluorescence microscopy after 2hrs, 6hrs, 24hrs, and 48hrs of initial cell seeding. Mean fluorescence intensity for every time point was quantified and compared between conditions.

### Induction of Fibroblasts

In order to obtain CAFs, 10ng/ml TGFβ (BioLegend) was used as a positive control to induce fibroblast differentiation. WI-38 cells were seeded onto coverslips with in serum-free medium for 6 hours (starvation). Then serum-free medium was replaced with WI-38 complete medium supplemented with 10ng/ml TGFβ. Cells were then cultured for 48 hours.

### Immunostaining and Fluorescence Imaging Staining for CAFs

Fibroblast cells were cultured under three conditions: cells on ccECM-containing coverslips, ccECM-free coverslips with TGFβ treatment, and ccECM-free coverslips without TGFβ treatment. After 48 hours, cells were fixed with 4% paraformaldehyde (PFA) and permeabilized with 0.1% Triton X-100 in PBS. Blocking was performed with 5% goat serum in PBS for 60 minutes at room temperature. Primary antibodies (FAP, Vimentin, FSP-1/s100A4, and PDGFRβ) from the CAF marker sampler kit (Cell Signaling Technologies) were applied at ratios of 1:200 and incubated overnight at +4°C. Secondary antibodies (Alexa Fluor 488-conjugated goat anti-rabbit), Alexa Fluor 647-conjugated phalloidin, and DAPI were applied at ratios of 1:200, 1:30, and 1:1000, respectively, for 40 minutes in the dark. The coverslips were then mounted and imaged using Leica SP8 Confocal Microscope.

### Staining for ccECM

MDA-MB-231 cells were seeded onto coverslips at 5×10^4^ cells/cm^2^ density and cultured for five days. Following decellularization by freeze-thaw cycle, coverslips containing ccECM were treated with 200U/ml DNAse to remove cell-free nuclei. The ccECM was then fixed with 4% PFA and blocked with 5% goat serum in PBS. Overnight incubation at +4°C was performed with primary antibodies: Laminin (Sigma) and Fibronectin (Sigma), both at a dilution of 1:200. Subsequently, secondary antibodies (Alexa Fluor 488 goat-anti-rabbit) at a dilution of 1:200, along with DAPI (1:1000), were applied for 40 minutes in the dark. The coverslips were mounted and imaged using Leica SP8 Confocal Microscope.

### Scanning Electron Microscopy (SEM)

5×10^4^ MDA-MB-231 cells/cm^2^ were grown on indium tin oxide coated coverslips (Teknoma Teknolojik Malzemeler) for 5 days. Then cells were removed using either the chemical extraction or freeze-thaw cycling. 3% PFA and 1.5% glutaraldehyde (GA) prefixation solution were used to fix decellularized coverslips for 2 hours. For SEM, decellularized coverslips were exposed to serial dehydration, applying an increasing concentration of ethyl alcohol (EtOH) at different time intervals (50% for 5 min, 70% for 10min, 80% for 10min, 90% for 15 min, and finally 100% for 20 min). After the serial dehydration process, coverslips were held in the desiccator until needed. The coverslips were imaged using Thermo Scientific Quanta 250 FEG scanning electron microscope.

### Image Analysis

#### Quantification of Immunostaining Results

Images were processed using ImageJ to calculate the fluorescence intensity of CAF markers. Images were first exposed to subtraction after removing the background using a Gaussian blur filter. Next, the measure command was used to calculate the protein concentration and normalize it to the number of cells as determined by the number of nuclei stained with DAPI.

### RNA Isolation and Q-RT PCR

Fibroblasts were flash-frozen after being cultured for 48 hours in the three different culture conditions. Total RNA extraction and cDNA production was performed using the PureLink® RNA Mini Kit (NucleospinRNA, #74095510) and the cDNA Synthesis Kit (ThermoScientifics, # K1622) according to the manufacturers’ instructions. The Roche LightCycler 96 Real-Time PCR Detection System performed PCR amplification and detection, utilizing the FastStart Essential DNA Green Master (Roche, 06402712001). The delta-delta Ct technique was utilized to calculate the relative amounts of mRNA and the relative mRNA values are provided as mean ± standard deviation of three independent experiments, normalized to control (glass in the absence of TGFβ) samples, with each gene’s expression data being normalized to the TATA-Box Binding Protein (TBP) gene. Primers are provided in supplementary table 1.

### Statistical Analysis

Unless otherwise noted, the significance of all data was assessed using a t-test (two-sample, assuming unequal variances). Reported n values are for biological repeats. Each biological repeat had at least 2 technical repeats. For imaging, at least 3 different field of views were captured for each technical repeat. The data on the charts were reported as mean ± standard error, and the statistical indicator was taken as * p < 0.05; ** p < 0.01; *** p < 0.005; **** p < 0.0001.

## Acknowledgments

The authors thank O. Yalcin-Ozuysal for critical reading of the manuscript, A. Eshete Mohammed for helpful discussions and Z. Sinem Yılmaz for access to scanning electron microscope at IZTECH Integrated Research Center. This work was supported by IZTECH Scientific Research Project Grant Number 2022IYTE-1-0071.

## CRediT authorship contribution statement

Eyup YONDEM: Conceptualization, Methodology, Investigation, Data analysis, Validation, Visualization, Writing - original draft, Writing - review & editing. Devrim Pesen-Okvur: Conceptualization, Methodology, Investigation, Validation, Data analysis, Visualization, Writing - original draft, Writing - review & editing, Supervision, Funding acquisition.

## REFERENCES

1. Virchow, R., Die Cellularpathologie in ihrer Begründung auf physiologische und pathologische Gewebelehre: zwanzig Vorlesungen, gehalten während der Monate Februar, März und April 1858 im pathologischen Institute zu Berlin. Hirschwald: 1859; Vol. 1.

2. Mueller, M. M.; Fusenig, N. E., Friends or foes—bipolar effects of the tumour stroma in cancer. Nature Reviews Cancer 2004, 4 (11), 839–849.

3. Lazard, D.; Sastre, X.; Frid, M. G.; Glukhova, M. A.; Thiery, J.-P.; Koteliansky, V., Expression of smooth muscle-specific proteins in myoepithelium and stromal myofibroblasts of normal and malignant human breast tissue. Proceedings of the National Academy of Sciences 1993, 90 (3), 999–1003.

4. De Wever, O.; Demetter, P.; Mareel, M.; Bracke, M., Stromal myofibroblasts are drivers of invasive cancer growth. International journal of cancer 2008, 123 (10), 2229–2238.

5. Calvo, F.; Ege, N.; Grande-Garcia, A.; Hooper, S.; Jenkins, R. P.; Chaudhry, S. I.; Harrington, K.; Williamson, P.; Moeendarbary, E.; Charras, G., Mechanotransduction and YAP-dependent matrix remodelling is required for the generation and maintenance of cancer-associated fibroblasts. Nature cell biology 2013, 15 (6), 637–646.

6. Cirri, P.; Chiarugi, P., Cancer associated fibroblasts: the dark side of the coin. American journal of cancer research 2011, 1 (4), 482.

7. Rønnov-Jessen, L.; Petersen, O., Induction of alpha-smooth muscle actin by transforming growth factor-beta 1 in quiescent human breast gland fibroblasts. Implications for myofibroblast generation in breast neoplasia. Laboratory investigation; a journal of technical methods and pathology 1993, 68 (6), 696–707.

8. Desmoulière, A.; Geinoz, A.; Gabbiani, F.; Gabbiani, G., Transforming growth factor-beta 1 induces alpha-smooth muscle actin expression in granulation tissue myofibroblasts and in quiescent and growing cultured fibroblasts. The Journal of cell biology 1993, 122 (1), 103–111.

9. William Petersen, O.; Lind Nielsen, H.; Gudjonsson, T.; Villadsen, R.; Rønnov-Jessen, L.; Bissell, M. J., The plasticity of human breast carcinoma cells is more than epithelial to mesenchymal conversion. Breast Cancer Research 2001, 3 (4), 1–5.

10. Zeisberg, E. M.; Potenta, S.; Xie, L.; Zeisberg, M.; Kalluri, R., Discovery of endothelial to mesenchymal transition as a source for carcinoma-associated fibroblasts. Cancer research 2007, 67 (21), 10123–10128.

11. Zeisberg, E. M.; Tarnavski, O.; Zeisberg, M.; Dorfman, A. L.; McMullen, J. R.; Gustafsson, E.; Chandraker, A.; Yuan, X.; Pu, W. T.; Roberts, A. B., Endothelial-to-mesenchymal transition contributes to cardiac fibrosis. Nature medicine 2007, 13 (8), 952–961.

12. Potenta, S.; Zeisberg, E.; Kalluri, R., The role of endothelial-to-mesenchymal transition in cancer progression. British journal of cancer 2008, 99 (9), 1375–1379.

13. Wessels, D. J.; Pradhan, N.; Park, Y. N.; Klepitsch, M. A.; Lusche, D. F.; Daniels, K. J.; Conway, K. D.; Voss, E. R.; Hegde, S. V.; Conway, T. P.; Soll, D. R., Reciprocal signaling and direct physical interactions between fibroblasts and breast cancer cells in a 3D environment. PLoS One 2019, 14 (6), e0218854.

14. Jeong, S.-Y.; Lee, J.-H.; Shin, Y.; Chung, S.; Kuh, H.-J., Co-culture of tumor spheroids and fibroblasts in a collagen matrix-incorporated microfluidic chip mimics reciprocal activation in solid tumor microenvironment. PloS one 2016, 11 (7), e0159013.

15. Jotzu, C.; Alt, E.; Welte, G.; Li, J.; Hennessy, B. T.; Devarajan, E.; Krishnappa, S.; Pinilla, S.; Droll, L.; Song, Y.-H., Adipose tissue derived stem cells differentiate into carcinoma-associated fibroblast-like cells under the influence of tumor derived factors. Cellular oncology 2011, 34 (1), 55–67.

16. Bissell, M. J.; Hall, H. G.; Parry, G., How does the extracellular matrix direct gene expression? Journal of theoretical biology 1982, 99 (1), 31–68.

17. Tian, C.; Clauser, K. R.; Öhlund, D.; Rickelt, S.; Huang, Y.; Gupta, M.; Mani, D.; Carr, S. A.; Tuveson, D. A.; Hynes, R. O., Proteomic analyses of ECM during pancreatic ductal adenocarcinoma progression reveal different contributions by tumor and stromal cells. Proceedings of the National Academy of Sciences 2019, 116 (39), 19609–19618.

18. Cox, T. R., The matrix in cancer. Nature Reviews Cancer 2021, 21 (4), 217–238.

19. Kim, S.-H.; Turnbull, J.; Guimond, S., Extracellular matrix and cell signalling: the dynamic cooperation of integrin, proteoglycan and growth factor receptor. The Journal of endocrinology 2011, 209 (2), 139–151.

20. Fattet, L.; Jung, H.-Y.; Matsumoto, M. W.; Aubol, B. E.; Kumar, A.; Adams, J. A.; Chen, A. C.; Sah, R. L.; Engler, A. J.; Pasquale, E. B., Matrix rigidity controls epithelial-mesenchymal plasticity and tumor metastasis via a mechanoresponsive EPHA2/LYN complex. Developmental cell 2020, 54 (3), 302–316. e7.

21. Hinderer, S.; Layland, S. L.; Schenke-Layland, K., ECM and ECM-like materials—Biomaterials for applications in regenerative medicine and cancer therapy. Advanced drug delivery reviews 2016, 97, 260–269.

22. Goh, S.-K.; Olsen, P.; Banerjee, I., Correction: Extracellular Matrix Aggregates from Differentiating Embryoid Bodies as a Scaffold to Support ESC Proliferation and Differentiation. Plos one 2013, 8 (10).

23. Tan, Q.; Xu, L.; Zhang, J.; Ning, L.; Jiang, Y.; He, T.; Luo, J.; Chen, J.; Lv, Q.; Yang, X., Breast cancer cells interact with tumor-derived extracellular matrix in a molecular subtype-specific manner. Biomaterials Advances 2023, 146, 213301.

24. Galindo-Pumarino, C.; Herrera, A.; Munoz, A.; Carrato, A.; Herrera, M.; Pena, C., Fibroblast-Derived 3D Matrix System Applicable to Endothelial Tube Formation Assay. J Vis Exp 2019, (154).

25. Liu, G.; Wang, B.; Li, S.; Jin, Q.; Dai, Y., Human breast cancer decellularized scaffolds promote epithelial-to-mesenchymal transitions and stemness of breast cancer cells in vitro. Journal of cellular physiology 2019, 234 (6), 9447–9456.

26. Franco-Barraza, J.; Beacham, D. A.; Amatangelo, M. D.; Cukierman, E., Preparation of extracellular matrices produced by cultured and primary fibroblasts. Current protocols in cell biology 2016, 71 (1), 10.9. 1–10.9. 34.

27. Cukierman, E., Preparation of extracellular matrices produced by cultured fibroblasts. Current protocols in cell biology 2002, 16 (1), 10.9. 1–10.9. 15.

28. Guido, C.; Whitaker-Menezes, D.; Capparelli, C.; Balliet, R.; Lin, Z.; Pestell, R. G.; Howell, A.; Aquila, S.; Ando, S.; Martinez-Outschoorn, U.; Sotgia, F.; Lisanti, M. P., Metabolic reprogramming of cancer-associated fibroblasts by TGF-beta drives tumor growth: connecting TGF-beta signaling with “Warburg-like” cancer metabolism and L-lactate production. Cell Cycle 2012, 11 (16), 3019–35.

29. Li, Q.; Zhang, D.; Wang, Y.; Sun, P.; Hou, X.; Larner, J.; Xiong, W.; Mi, J., MiR-21/Smad 7 signaling determines TGF-beta1-induced CAF formation. Sci Rep 2013, 3, 2038.

30. Bordignon, P.; Bottoni, G.; Xu, X.; Popescu, A. S.; Truan, Z.; Guenova, E.; Kofler, L.; Jafari, P.; Ostano, P.; Rocken, M.; Neel, V.; Dotto, G. P., Dualism of FGF and TGF-beta Signaling in Heterogeneous Cancer-Associated Fibroblast Activation with ETV1 as a Critical Determinant. Cell Rep 2019, 28 (9), 2358–2372 e6.

31. Yoon, H.; Tang, C. M.; Banerjee, S.; Delgado, A. L.; Yebra, M.; Davis, J.; Sicklick, J. K., TGF-beta1-mediated transition of resident fibroblasts to cancer-associated fibroblasts promotes cancer metastasis in gastrointestinal stromal tumor. Oncogenesis 2021, 10 (2), 13.

32. Han, C.; Liu, T.; Yin, R., Biomarkers for cancer-associated fibroblasts. Biomark Res 2020, 8 (1), 64.

33. Sahai, E.; Astsaturov, I.; Cukierman, E.; DeNardo, D. G.; Egeblad, M.; Evans, R. M.; Fearon, D.; Greten, F. R.; Hingorani, S. R.; Hunter, T.; Hynes, R. O.; Jain, R. K.; Janowitz, T.; Jorgensen, C.; Kimmelman, A. C.; Kolonin, M. G.; Maki, R. G.; Powers, R. S.; Pure, E.; Ramirez, D. C.; Scherz-Shouval, R.; Sherman, M. H.; Stewart, S.; Tlsty, T. D.; Tuveson, D. A.; Watt, F. M.; Weaver, V.; Weeraratna, A. T.; Werb, Z., A framework for advancing our understanding of cancer-associated fibroblasts. Nat Rev Cancer 2020, 20 (3), 174–186.

34. Ostrowska-Podhorodecka, Z.; Ding, I.; Norouzi, M.; McCulloch, C. A., Impact of Vimentin on Regulation of Cell Signaling and Matrix Remodeling. Frontiers in Cell and Developmental Biology 2022, 10, 869069–869069.

35. Sarrio, D.; Rodriguez-Pinilla, S. M.; Hardisson, D.; Cano, A.; Moreno-Bueno, G.; Palacios, J., Epithelial-mesenchymal transition in breast cancer relates to the basal-like phenotype. Cancer Res 2008, 68 (4), 989–97.

36. Vuoriluoto, K.; Haugen, H.; Kiviluoto, S.; Mpindi, J. P.; Nevo, J.; Gjerdrum, C.; Tiron, C.; Lorens, J. B.; Ivaska, J., Vimentin regulates EMT induction by Slug and oncogenic H-Ras and migration by governing Axl expression in breast cancer. Oncogene 2011, 30 (12), 1436–48.

37. Kelly, T.; Huang, Y.; Simms, A. E.; Mazur, A., Fibroblast activation protein-α: a key modulator of the microenvironment in multiple pathologies. International review of cell and molecular biology 2012, 297, 83–116.

38. Donovan, J.; Shiwen, X.; Norman, J.; Abraham, D., Platelet-derived growth factor alpha and beta receptors have overlapping functional activities towards fibroblasts. Fibrogenesis & tissue repair 2013, 6 (1), 1–9.

39. Österreicher, C. H.; Penz-Österreicher, M.; Grivennikov, S. I.; Guma, M.; Koltsova, E. K.; Datz, C.; Sasik, R.; Hardiman, G.; Karin, M.; Brenner, D. A., Fibroblast-specific protein 1 identifies an inflammatory subpopulation of macrophages in the liver. Proceedings of the National Academy of Sciences 2011, 108 (1), 308–313.

40. Verstegen, M. M.; Willemse, J.; Van Den Hoek, S.; Kremers, G.-J.; Luider, T. M.; van Huizen, N. A.; Willemssen, F. E.; Metselaar, H. J.; IJzermans, J. N.; van der Laan, L. J., Decellularization of whole human liver grafts using controlled perfusion for transplantable organ bioscaffolds. Stem cells and development 2017, 26 (18), 1304–1315.

41. Hielscher, A. C.; Qiu, C.; Gerecht, S., Breast cancer cell-derived matrix supports vascular morphogenesis. American Journal of Physiology-Cell Physiology 2012, 302 (8), C1243–C1256.

42. Mhaidly, R.; Mechta-Grigoriou, F., Role of cancer-associated fibroblast subpopulations in immune infiltration, as a new means of treatment in cancer. Immunological reviews 2021, 302 (1), 259–272.

43. Nurmik, M.; Ullmann, P.; Rodriguez, F.; Haan, S.; Letellier, E., In search of definitions: Cancer-associated fibroblasts and their markers. Int J Cancer 2020, 146 (4), 895–905.

44. Friedman, G.; Levi-Galibov, O.; David, E.; Bornstein, C.; Giladi, A.; Dadiani, M.; Mayo, A.; Halperin, C.; Pevsner-Fischer, M.; Lavon, H.; Mayer, S.; Nevo, R.; Stein, Y.; Balint-Lahat, N.; Barshack, I.; Ali, H. R.; Caldas, C.; Nili-Gal-Yam, E.; Alon, U.; Amit, I.; Scherz-Shouval, R., Cancer-associated fibroblast compositions change with breast cancer progression linking the ratio of S100A4(+) and PDPN(+) CAFs to clinical outcome. Nat Cancer 2020, 1 (7), 692–708.

45. Zhao, Z.; Li, T.; Yuan, Y.; Zhu, Y., What is new in cancer-associated fibroblast biomarkers? Cell Communication and Signaling 2023, 21 (1), 1–23.

46. Zhang, X.; Chen, X.; Hong, H.; Hu, R.; Liu, J.; Liu, C., Decellularized extracellular matrix scaffolds: Recent trends and emerging strategies in tissue engineering. Bioact Mater 2022, 10, 15–31.

47. Assuncao, M.; Dehghan-Baniani, D.; Yiu, C. H. K.; Spater, T.; Beyer, S.; Blocki, A., Cell-Derived Extracellular Matrix for Tissue Engineering and Regenerative Medicine. Front Bioeng Biotechnol 2020, 8, 602009.

48. Winkler, J.; Abisoye-Ogunniyan, A.; Metcalf, K. J.; Werb, Z., Concepts of extracellular matrix remodelling in tumour progression and metastasis. Nature communications 2020, 11 (1), 5120.

49. Xiong, G.-F.; Xu, R., Function of cancer cell-derived extracellular matrix in tumor progression. Journal of Cancer Metastasis and Treatment 2016, 2 (9).

50. Poltavets, V.; Kochetkova, M.; Pitson, S. M.; Samuel, M. S., The Role of the Extracellular Matrix and Its Molecular and Cellular Regulators in Cancer Cell Plasticity. Front Oncol 2018, 8, 431.

51. Vogel, C.; Marcotte, E. M., Insights into the regulation of protein abundance from proteomic and transcriptomic analyses. Nature reviews genetics 2012, 13 (4), 227–232.

52. Ba, P.; Zhang, X.; Yu, M.; Li, L.; Duan, X.; Wang, M.; Lv, S.; Fu, G.; Yang, P.; Yang, C.; Sun, Q., Cancer associated fibroblasts are distinguishable from peri-tumor fibroblasts by biological characteristics in TSCC. Oncol Lett 2019, 18 (3), 2484–2490.

53. Yu, T.; Guo, Z.; Fan, H.; Song, J.; Liu, Y.; Gao, Z.; Wang, Q., Cancer-associated fibroblasts promote non-small cell lung cancer cell invasion by upregulation of glucose-regulated protein 78 (GRP78) expression in an integrated bionic microfluidic device. Oncotarget 2016, 7 (18), 25593.

54. Nomura, S., Identification, friend or foe: vimentin and α-smooth muscle actin in cancer-associated fibroblasts. Annals of Surgical Oncology 2019, 26 (13), 4191–4192.

55. Casey, T. M.; Eneman, J.; Crocker, A.; White, J.; Tessitore, J.; Stanley, M.; Harlow, S.; Bunn, J. Y.; Weaver, D.; Muss, H.; Plaut, K., Cancer associated fibroblasts stimulated by transforming growth factor beta1 (TGF-beta 1) increase invasion rate of tumor cells: a population study. Breast Cancer Res Treat 2008, 110 (1), 39–49.

56. Lugo-Cintrón, K. M.; Gong, M. M.; Ayuso, J. M.; Tomko, L. A.; Beebe, D. J.; Virumbrales-Muñoz, M.; Ponik, S. M., Breast fibroblasts and ECM components modulate breast cancer cell migration through the secretion of MMPs in a 3D microfluidic co-culture model. Cancers 2020, 12 (5), 1173.

