## Supplemental Data for "Cancer cell-derived extracellular matrix promotes differentiation of fibroblasts into cancer-associated fibroblasts"

**Supp. Figure 1.** Coomassie blue staining was used to detect the presence of proteins on coverslips.

Coomassie blue staining for coverslips coated with different proteins at different concentrations. 0.1mg/ml Matrigel, 0.05mg/ml collagen, 0.1mg/ml poly-l-lysine (PLL). A culture medium without FBS(serum-free) coated coverslip was used as a control.

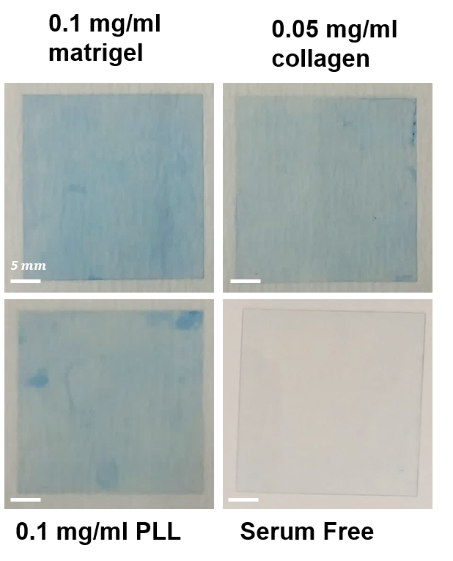

**Supp. Fıgure 2.** DNAse-1 treatment at 200 U/mL on decellularized coverslips was sufficient for eliminating cell-free nuclei prior to analysis.

DNAse-1 treatment at 200 U/mL on decellularized coverslips was sufficient for eliminating cell-free nuclei prior to analysis. A. Fluorescence and phase contrast images. MDA-MB-231 cells (cc) in culture, ccECM without treatment, ccECM with 100 U/ml DNAse treatment, ccECM with 200 U/ml DNAse treatment. Cells were cultured for 5 days.

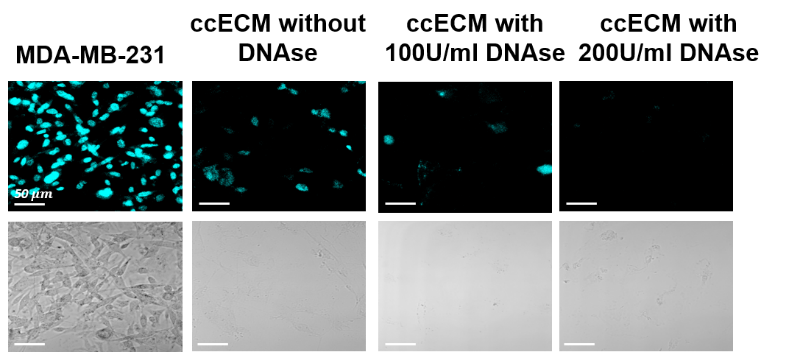

**Supp. Table 1.** Forward and reverse primers were used in this study to reveal CAFs’ marker expression.

| Genes | Forward (5’---3^’^) Primers | Reverse (5’---3^’^) Primers |
| --- | --- | --- |
| TBP | TAGAAGGCCTTGTGCTCACC | TCTGCTCTGACTTTAGCACCT |
| Vimentin | GCTAACCAACGACAAAGCCC | CGTTCAAGGTCAAGACGTGC |
| ACTA2/α-SMA | TCAATGTCCCAGCCATGTAT | CAGCACGATGCCAGTTGT |
| FAP | GAAAGAAAGGTGCCAATA | GATCAGTGCGTCCATCA |
| PDGFRβ | ACA CGG GAG AAT ACT TTT GC | GTT CCT CGG CAT CAT TAG GG |
| FSP-1/s100A4 | TCCACAAGTACTCGGGCAAAG | CTCTTGGAAGTCCACCTCGT |
